# National Database of Health Insurance Claims and Specific Health Checkups of Japan (NDB): Outline and Patient-Matching Technique

**DOI:** 10.1101/280008

**Authors:** Kubo Shinichiro, Noda Tatsuya, Myojin Tomoya, Nishioka Yuichi, Higashino Tsuneyuki, Matsui Hiroki, Kato Genta, Imamura Tomoaki

## Abstract

**Background:** The National Database of Health Insurance Claims and Specific Health Checkups of Japan (NDB) is a comprehensive database of health insurance claims data under Japan’s National Health Insurance system. The NDB uses two types of personal identification variables (referred to in the database as “ID1” and “ID2”) to link the insurance claims of individual patients. However, the information entered against these ID variables is prone to change for several reasons, such as when claimants find or change employment, or due to variations in the spelling of their name. In the present study, we developed a new patient-matching technique that improves upon the existing system of using ID1 and ID2 variables. We also sought to validate a new personal ID variable (ID0) that we propose in order to enhance the efficiency of patient matching in the NDB database.

**Methods:** Our study targeted data from health insurance claims filed between April 2013 and March 2016 for hospitalization, combined diagnostic procedures, outpatient treatment, and dispensing of prescription medication. We developed a new patient-matching algorithm based on the ID1 and ID2 variables, as well as variables for treatment date and clinical outcome. We then attempted to validate our algorithm by comparing the number of patients identified by patient matching with the current ID1 variable and our proposed ID0 variable against the estimated patient population as of 1 October 2015.

**Results:** The numbers of patients in each sex and age group that were identified with the ID0 variable were lower than those identified using the ID1 variable. By using the ID0 variable, we were able to reduce the number of duplicate records for male and female patients by 5.8% and 6.4%, respectively. The numbers of children, adults older than 75 years, and women of reproductive age identified using the ID1 patient-matching variable were all higher than their corresponding estimates. Conversely, the numbers of these patients identified with the ID0 patient-matching variable were all within their corresponding estimates.

**Conclusion:** Our findings show that the proposed ID0 variable delivers more precise patient-matching results than the existing ID1 variable. The ID0 variable is currently the best available technique for patient matching in the NDB database. Future patient population estimates should therefore rely on the ID0 variable instead of the ID1 variable.

## 1. Introduction

Universal health coverage was established in Japan in 1961. Health service costs are largely covered by contributions from insurance societies or public funding, with copayment (up to 30% of the cost) that varies depending on age and income of beneficiaries and types of medical services provided. Hospitals and clinics prepare health insurance claims for individual patients every month and send them to the corresponding insurers through the Health Insurance Claims Review and Reimbursement Services, thereby receiving reimbursement accordingly. In 2006, a national database managed by the Japan Ministry of Health, Labour and Welfare containing data of health insurance claims and data of specific health checkups, was established to serve as a valuable resource for obtaining statistical data useful in formulating plans regarding medical care expenditure regulations^1^. Supported by the promotion of issuing health insurance claims online or via electronic media, the national database of health insurance claims and specific health checkups (NDB) was created to store anonymized data included in health insurance claims issued in April 2009 and beyond. The NDB stores dental claims data and data regarding specific health checkups and specific health guidance given to beneficiaries. The specific health checkups and specific health guidance program, targeting insured and dependents aged 40-74 years, is one of several measures to prevent life-style diseases. Thus, the NDB stores information about findings from history taking, test results, and health guidance contents.

Given that Japan has a universal health coverage system, the NDB is one of the world’s largest health-related databases and contains complete datasets of insured medical care, including information on approximately 12,884 million claims from over a hundred million individuals issued between April 2009 and December 2016 (data as of March 2017)^2^. The contents of the NDB are findings from examinations by doctors, but not judgements by other professionals or patients themselves. Furthermore, health insurance claims are prepared and submitted to be reviewed for reimbursement, and thus they include valid information regarding medical care provided and drugs dispensed. Thus, effective use of the NDB enables retrospective cohort studies with a sample size of around 100 million, and other types of studies, with very small selection bias.

However, use of the NDB is currently limited. According to our previous study, this is mainly because the volume of information contained is very large, and the format of the insurance claims, originally designed for the reimbursement process, is not suited for research without modification^3^. In particular, identifying, tracking, and linking data corresponding to a given patient is a major challenge.

Health insurance claims are prepared for individual patients every month. A single patient often seeks medical care over multiple months, or visits different medical institutions within the same month, and thus time-course analysis cannot be appropriately conducted without linking data corresponding to a given individual. Such data linkage is supposed to be possible with ID1 and ID2, which are anonymized variables shown as sequences of alphanumeric characters. ID1 is a hash value generated from the insurer’s ID, and the beneficiary’s ID, date of birth, and sex; ID2 is a hash value generated from the beneficiary’s name, date of birth, and sex. However, ID1 is not permanent, because the insurer can change when the beneficiary becomes employed or changes jobs. Similarly, ID2 can change when the beneficiary gets married or divorced, and due to orthographic variation among different medical institutions (e.g. use of different Chinese characters: 渡辺 vs 渡邉, which are both read as “Watanabe”^4^). This means that these IDs do not link data corresponding to the same person before and after a life event (e.g. finding employment, career change), but wrongly indicate them as belonging to different individuals. This may result in considerable errors in estimating the number of patients and in any estimates on a per patient basis.

There are two types of errors in linking data of health insurance claims corresponding to a given patient: wrongly linking health insurance claims that belong to different individuals (Type I error, Figure 1) and not linking health insurance claims belonging to the same individual (Type II error, Figure 2).

**Figure 1.**
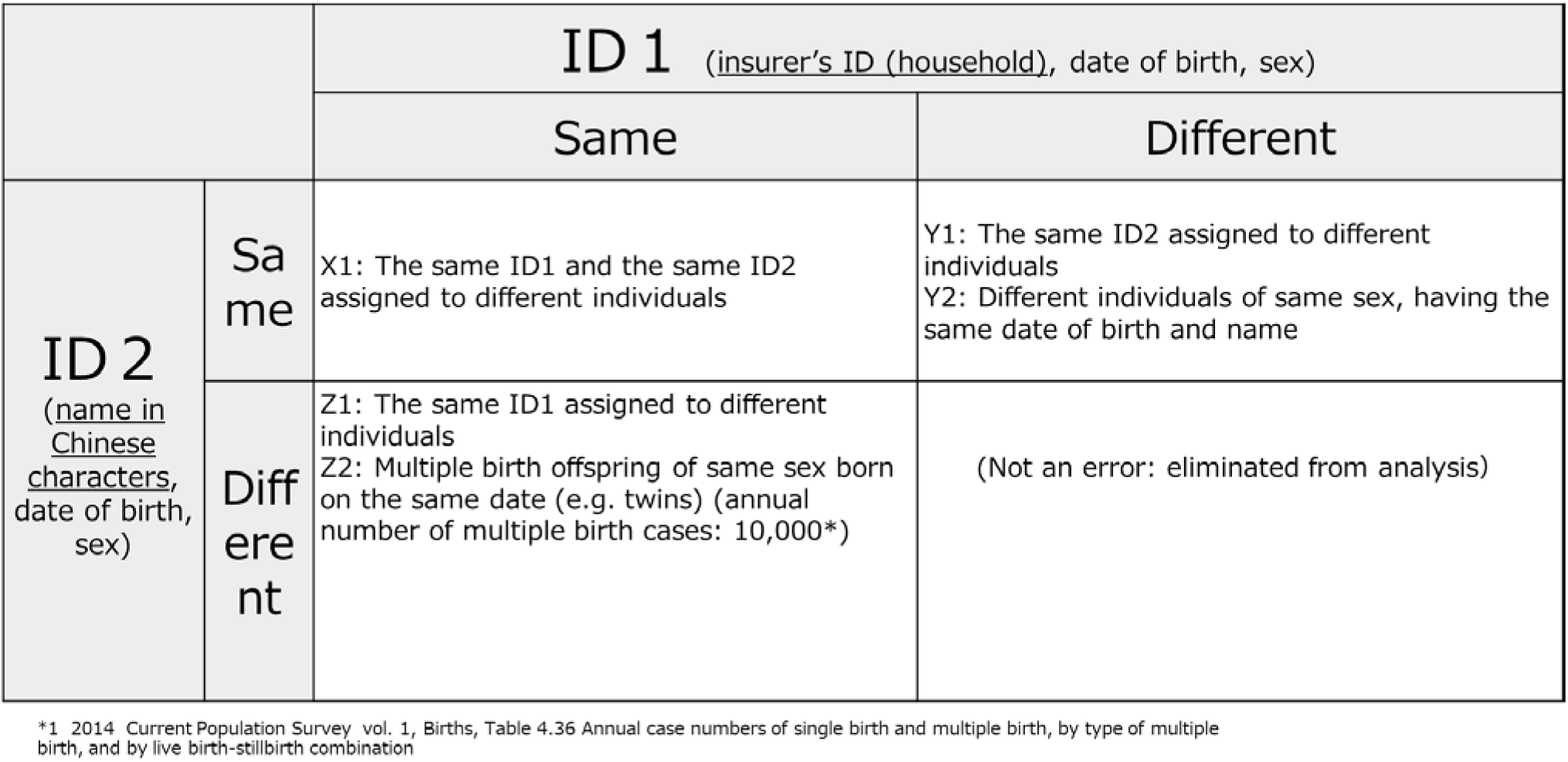
Assignment of the same ID to different individuals (type I error)

**Figure 2:**
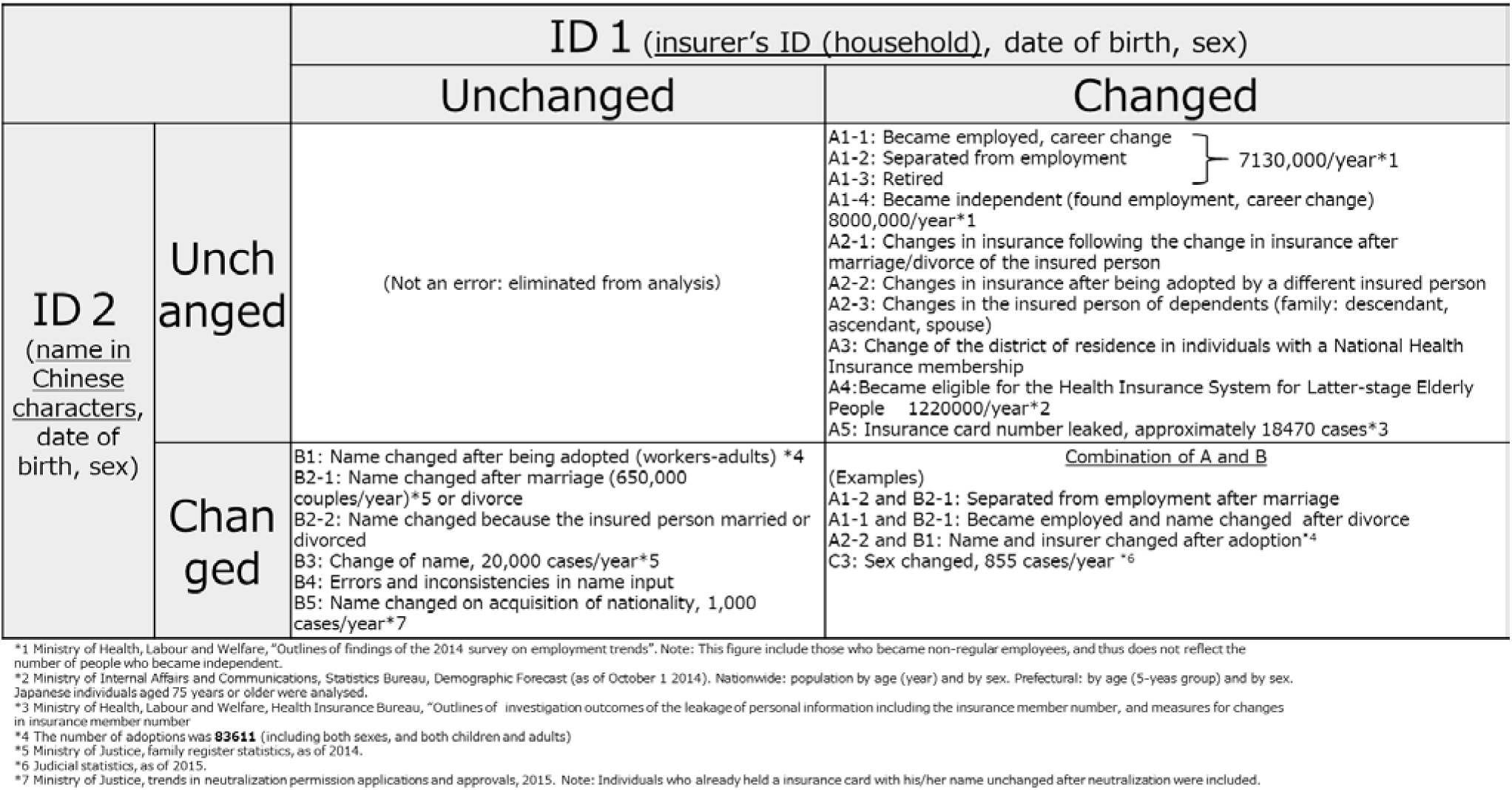
Assignment of different IDs to the same individual (type II error)

The same ID2 can be assigned to different individuals (type I error) when their names, dates of birth and sex are identical. People with the same family name and given name are not rare, and certain family names are very common in certain regions (e.g. Okinawa Prefecture). Also, there are trends in given names depending on the birth year. Thus, the number of individuals of same sex with the same name, and date of birth is not negligible.

The ID1 of the same patient changes (type II error) when the insurer’s ID, and consequently, beneficiary’s ID change. This occurs following a life event, such as career change, loss of employment, and retirement. The impact of change in insurer is substantial, because it also affects ID1s of dependents of the insured individual. Also, ID1 of a national health insurance changes each time the insured individual relocates. Furthermore, everybody’s ID1 changes at age 75 years when the Health Insurance System for Latter-stage Elderly People takes effect. Taken together, a completely different ID1 can be given to the same patient after a life event. Meanwhile, ID2 changes mainly due to change of name (change of family name upon adoption or marriage), and errors and inconsistencies in name input such as (1) differences in characters used (e.g. all Chinese character vs all Katakana; 鈴木太郎 vs スズキタロウ for “Suzuki Taro”); (2) difference in use of space between family and given names (e.g. 鈴木太郎 vs 鈴木太郎); (3) difference in character encoding such as use of full- and half-width characters (e.g. スズキタロウ vs ｽｽﾞｷﾀﾛｳ); and (4) difference in glyph used (e.g. 渡辺 vs 渡邉). It is very difficult to link data if ID1 and ID2 change simultaneously. For example, both name and the insurer change in roughly synchronized timing when an insured person retires from their job upon marriage or adoption.

To address these problems, a unique personal ID that remains unchanged throughout life needs to be used in the NDB. Such an ID system is likely to be fully available in the field of health care after 2020. This unique personal ID system is not beneficial in analyzing existing data and use of ID1 and ID2 are the only option. Meanwhile, use of ID3 was proposed for better data linkage^5^. ID3 eliminates orthographic variations affecting ID1 (full- or half-width, with/without an additional zero in front) for better linkage between data of specific health checkups and data of health insurance claims. Thus, the process of linking different IDs possibly corresponding to the same individual is still required.

The objectives of this study are to propose a new personal ID (ID0) that achieves, with various modifications, more efficient data linkage than the ones currently available, and to verify the new ID0-based approach.

## 2. Methods

Subjects of this study were medical inpatient claims, medical outpatient claims, diagnosis procedure combination (DPC) claims, and pharmacy claims, but not dental claims and specific health checkups data, issued in a 36-month period (April 2013-March 2016). DPC claims were defined as claims corresponding to bundled payments during DPC hospitalization at a DPC institution (claims for non-bundled payments were categorized as inpatient claims). A new algorithm using ID1, ID2, data of medical care received, and medical care outcome was tested to track changes in IDs of the same individual over multiple months. Given that medicine prescribed for outpatients is likely to be dispensed at pharmacies outside the medical institution, one-to-one linkage would become more difficult if pharmacy claims data are included in en bloc data processing. Thus, an intermediate data set was prepared from DPC and medical care (inpatient and outpatient) claims, and a separate intermediate data set was prepared from pharmacy claims data, and the two were examined side by side to obtain complete linkage. The following is the newly developed algorithm for linking data corresponding to a given individual.

### 2.1 Preparation of an intermediate data set from medical (inpatient and outpatient) and DPC claims

Figure 3 shows an example of an intermediate data set. ID1, ID2, date of medical care, and medical care outcomes were extracted from “medical inpatient claims”, “medical outpatient claims”, “medical inpatient claims subjected to bundled payment during DPC hospitalization”, “DPC claims during DPC hospitalization” and “DPC claims subjected to bundled payment during DPC hospitalization”. First, data dated within a period of a few months were searched for the same ID1, which were considered to correspond to the same individual. The data linkage process ended when death was noted in the outcome section.

**Figure 3.**
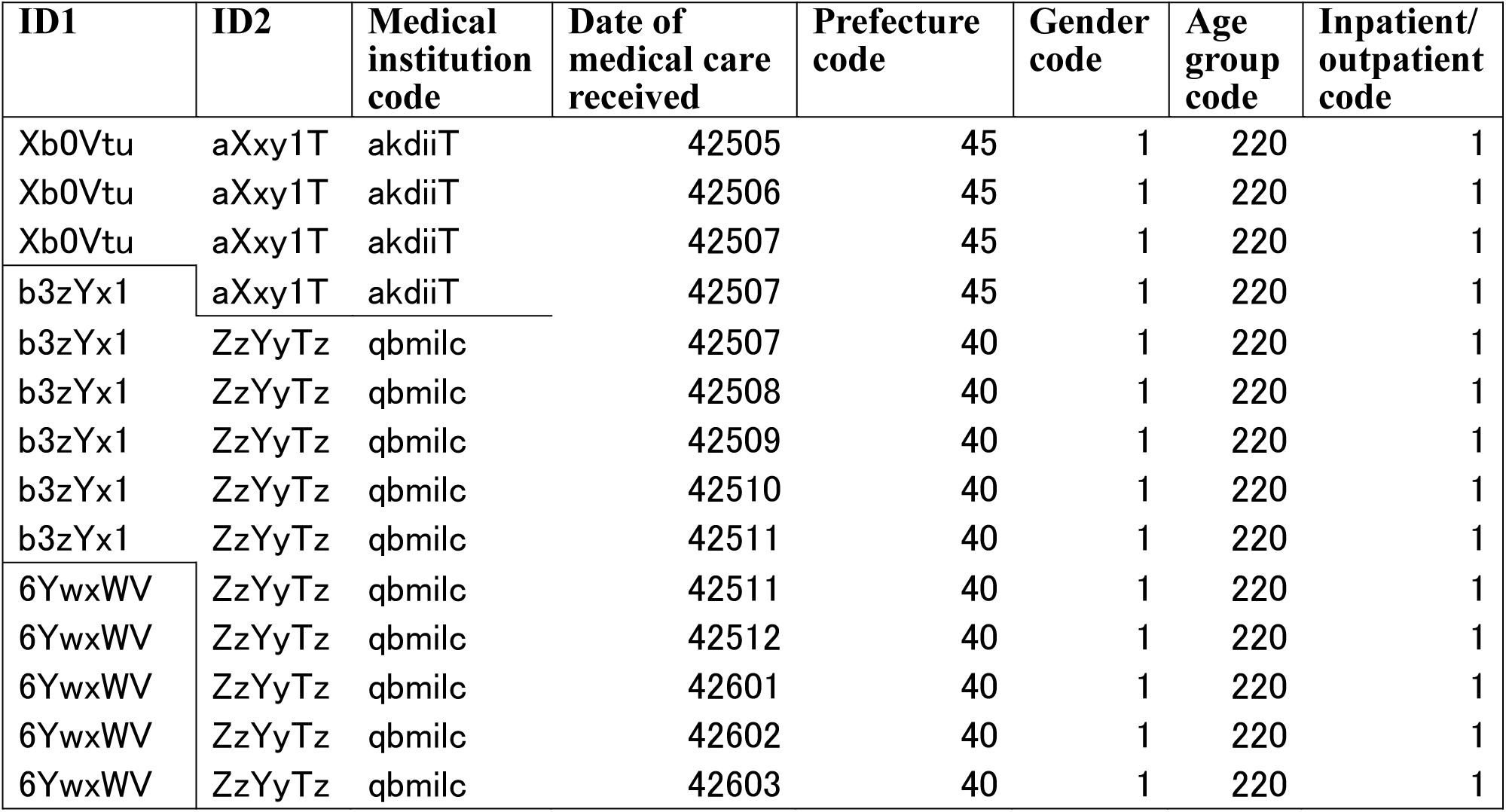
Example of data linking

When a sequence of claims tracked by an ID1 ended at a certain time, the corresponding ID2 around that timing was used for further tracking. When multiple ID2 candidates were found, the data linkage process was ended to avoid linkage of data corresponding to different individuals.

Claims associated with an individual whose eligibility is under question need to be re-reviewed by the insurer through the Health Insurance Claims Review and Reimbursement Services. Such claims for re-review are not included in the NDB, and the old ID1 and a newly assigned ID1 can co-exist for about 3 months. To address this problem, the ID2 corresponding to the old ID1 was used for search data in the following month, in the first preceding month, and then in the second preceding month to obtain an intermediate dataset table (medical and DPC).

### 2.2 Preparation of an intermediate dataset from pharmacy claims

ID1, ID2, and date of medical care were extracted from pharmacy claims. Data dated within a period of a few months were searched for the same ID1, and were considered to correspond to the same individual. When a sequence of claims with ID1 ended, ID2 was used for further tracking in a similar manner to that described in 2.1.

### 2.3 Linkage of medical claims (inpatient and outpatient) and pharmacy claims

Two types of intermediate dataset tables (see above) were linked to make a table that chronologically paired different ID1s belonging to the same individual (one-to-one ID1 pairing table). The left ID1 and the right ID1, assigned to the same person, in the same row are henceforth referred to as the old ID1 and the new ID1, respectively (e.g. p8d89jss is the old ID1 and ue8k22ue is the new ID1 in the p8d89jss-ue8k22ue pair). The old ID1 is the first ID1 issued irrespective of the category of the claim. When a set of reversible pairs (the kwyrls5T-Loi2g7Zx pair and the Loi2g7Zx-kwyrls5T pair) is observed, one pair is eliminated from the analysis.

### 2.4 Linking data corresponding to a given individual

Using the initial data-linking table, the new ID1 was replaced by the old ID1 within a row so that a single ID1 was assigned (provisional ID0). When the replaced ID1 was the old ID1 of the different pair, it was replaced by the new ID1. This process was repeated until all second ID1s were replaced. The remaining ID1 was defined as a new variable ID0 (Figure 4). In other words, by replacing an old ID1 sequentially with a new ID1, a new data-linkage variable ID0 was obtained. When there were no linkable claims because only one visit was made to seek medical care (hence no one-to-one ID1 pairing), ID1 of the single claim served as ID0.

**Figure 4.**
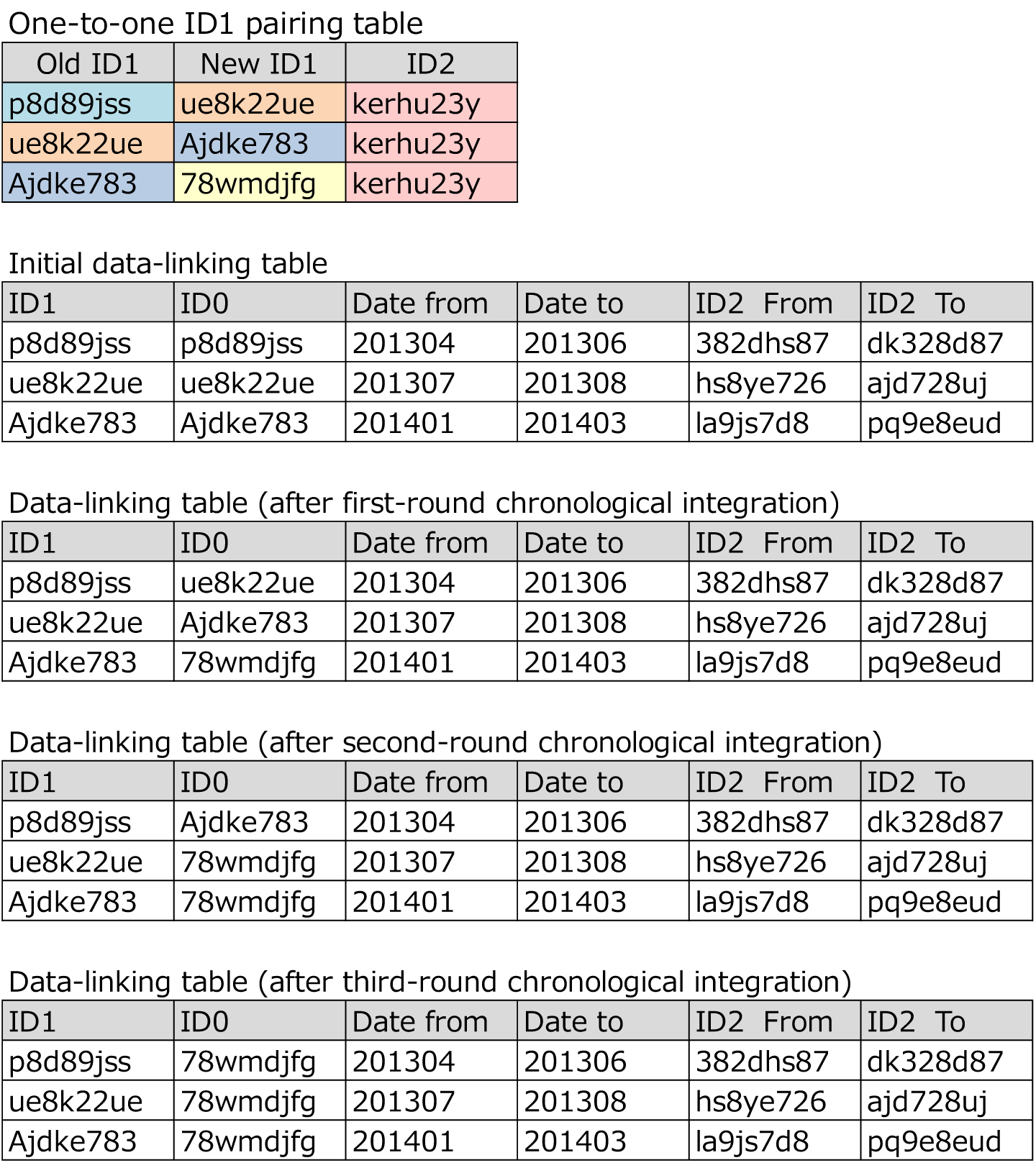
ID-linking process

The number of patients of each sex in each age group was estimated using ID0-based data aggregation, and resulting estimates were compared with those from ID1-based data aggregation in order to confirm the validity of this new variable. In addition, the rates of beneficiaries in the NDB (henceforth, the estimated rate of patients) were calculated using the estimated population (as of October 1, 2015), published by the Statistics Bureau, Ministry of Internal Affairs and Communications.

This study was approved by the Nara Medical School Ethics Committee (October 8, 2015, approval number 1123) and conducted in compliance with the Ethical Guidelines for Medical and Health Research Involving Human Subjects (announced in 2014, by the Ministry of Education, Culture, Sports, Science and Technology, and the Ministry of Health, Labour and Welfare).

## 3. Results

### 3.1 Characteristics of ID1 and ID2 in the NDB

There are three ID1-ID2 combinations in terms of changes over time: (1) both unchanged; (2) one is changed; and (3) both are changed. The 1:1 pairing was maintained in the case of (1) above, while multiple ID1s or ID2s could be linked to a single ID2 or ID1, respectively in the cases of (2) and (3) above. For example, change of job and resulting change of insurer without change of name results in two ID2s corresponding to one ID1 in the NDB. Table 1 shows the occurrence rates of multiple ID1s or ID2s corresponding to a single ID2 or ID1, respectively, in the 36-month period.

**Table 1.** Multiple ID1s and ID2s linking to a single ID2 and ID1, respectively

| Claims | Multiple ID2s to a single ID1 |  |  | Multiple ID1s to a single ID2 |  |  |
| --- | --- | --- | --- | --- | --- | --- |
|  | No. of IDs | No. of case | rate vs total | No. of ID | No. of case | rate vs total |
| DPC | 1 | 6,551,879 | 98.60% | 1 | 6,598,453 | 98.90% |
|  | 2 | 95,502 | 1.40% | 2 | 71,336 | 1.10% |
|  | 3 | 500 | 0.00% | 3 | 1,125 | 0.00% |
|  | 4 | +++ | 0.00% | 4 | 23 | 0.00% |
|  | 5 | *** | 0.00% |  |  |  |
|  | 6 | *** | 0.00% |  |  |  |
|  | Total | 6,647,932 | 100% |  | 6,670,937 | 100% |
| Medical inpatient | 1 | 5,600,307 | 98.00% | 1 | 5,661,120 | 98.60% |
|  | 2 | 112,202 | 2.00% | 2 | 79,723 | 1.40% |
|  | 3 | 974 | 0.00% | 3 | 2,221 | 0.00% |
|  | 4 | 17 | 0.00% | 4 | +++ | 0.00% |
|  |  |  |  | 5 | *** | 0.00% |
|  | Total | 5,713,500 | 100% |  | 5,743,180 | 100% |
| Medical outpatient | 1 | 93,145,865 | 79.60% | 1 | 122,674,241 | 92.90% |
|  | 2 | 22,651,589 | 19.40% | 2 | 8,568,817 | 6.50% |
|  | 3 | 1,141,050 | 1.00% | 3 | 726,050 | 0.50% |
|  | 4 | 96,966 | 0.10% | 4 | 71,728 | 0.10% |
|  | 5 | 9,678 | 0.00% | 5 | 8,055 | 0.00% |
|  | 6 | 2,192 | 0.00% | 6 | 1,250 | 0.00% |
|  | 7 | 527 | 0.00% | 7 | 222 | 0.00% |
|  | 8 | 155 | 0.00% | 8 | 60 | 0.00% |
|  | 9 | 54 | 0.00% | 9 | 29 | 0.00% |
|  | 10 | 16 | 0.00% | 10 | 15 | 0.00% |
|  | 11 | *** | 0.00% | 11 | *** | 0.00% |
|  | 12 | *** | 0.00% | 12 | *** | 0.00% |
|  | 14 | *** | 0.00% |  |  |  |
|  | Total | 117,048,101 | 100% |  | 132,050,478 | 100% |
| Pharmacy | 1 | 87,623,580 | 92.30% | 1 | 89,656,685 | 93.50% |
|  | 2 | 7,006,675 | 7.40% | 2 | 5,707,316 | 6.00% |
|  | 3 | 282,877 | 0.30% | 3 | 439,799 | 0.50% |
|  | 4 | 21,838 | 0.00% | 4 | 41,828 | 0.00% |
|  | 5 | 2,073 | 0.00% | 5 | 4,623 | 0.00% |
|  | 6 | 474 | 0.00% | 6 | 714 | 0.00% |
|  | 7 | 87 | 0.00% | 7 | 140 | 0.00% |
|  | 8 | 16 | 0.00% | 8 | 42 | 0.00% |
|  | 9 | *** | 0.00% | 9 | 14 | 0.00% |
|  | 10 | *** | 0.00% | 10 | *** | 0.00% |
|  | 14 | *** | 0.00% | 11 | *** | 0.00% |
|  |  |  |  | 12 | *** | 0.00% |
|  | Total | 94,937,633 | 100% |  | 95,851,172 | 100% |

\*\*\*: not disclosed due to a small number (<10), +++: after masking (≥10)

The rates of ID1s linked to only one ID2 (one-to-one pairing) were high (≥98%) in DPC claims and medical inpatient claims, but were relatively low in medical outpatient claims (79.6%) and pharmacy claims (92.3%). Such one-to-one pairing can be due to a single claim issued during the study period, or simultaneous changes in both IDs on a single claim (appearing as the first claim of an independent individual, although it is one of multiple claims corresponding to the same given individual), and all such claims were counted in this study.

The rate of ID2s linked to only one ID1 was 92.9% in medical outpatient claims, while that of ID1s linked to only one ID2 dropped markedly to 79.6%. This may be explained by orthographic variations of names that appear to occur frequently if patients visit multiple medical institutions. This influences the quality of linking of data corresponding to a given beneficiary.

### 3.2 Comparison of the accuracy between ID0-based and ID1-based data linking

FY2015 data, including PDC, medical inpatient, medical outpatient, and pharmacy claims, were analyzed using either ID0 or ID1 to estimate the number of patients by age group. The proportion of beneficiaries in the NDB (estimated rate of patients) to the estimated population (as of October 2015) by age group was also calculated (Tables 2 and 3). The numbers of both male and female patients were smaller when data was linked with ID0 than with ID1. The rate of the number of newly linked claims by use of ID0 to the number of patients by ID1-based linkage was 5.8% in men and 6.4% in women. The numbers of male and female patients aged 0-9, 65-59, and ≥75 years, and of female patients aged 20-35 years by ID1-based linkage markedly exceeded the corresponding estimated populations. In contrast, the numbers of male and female patients aged 0-9, male patients aged ≥80 years, and female patients ≥85 years by ID0-based data linkage markedly exceeded the corresponding estimated populations, but those of patients in other age groups were within the estimated populations.

**Table 2.**
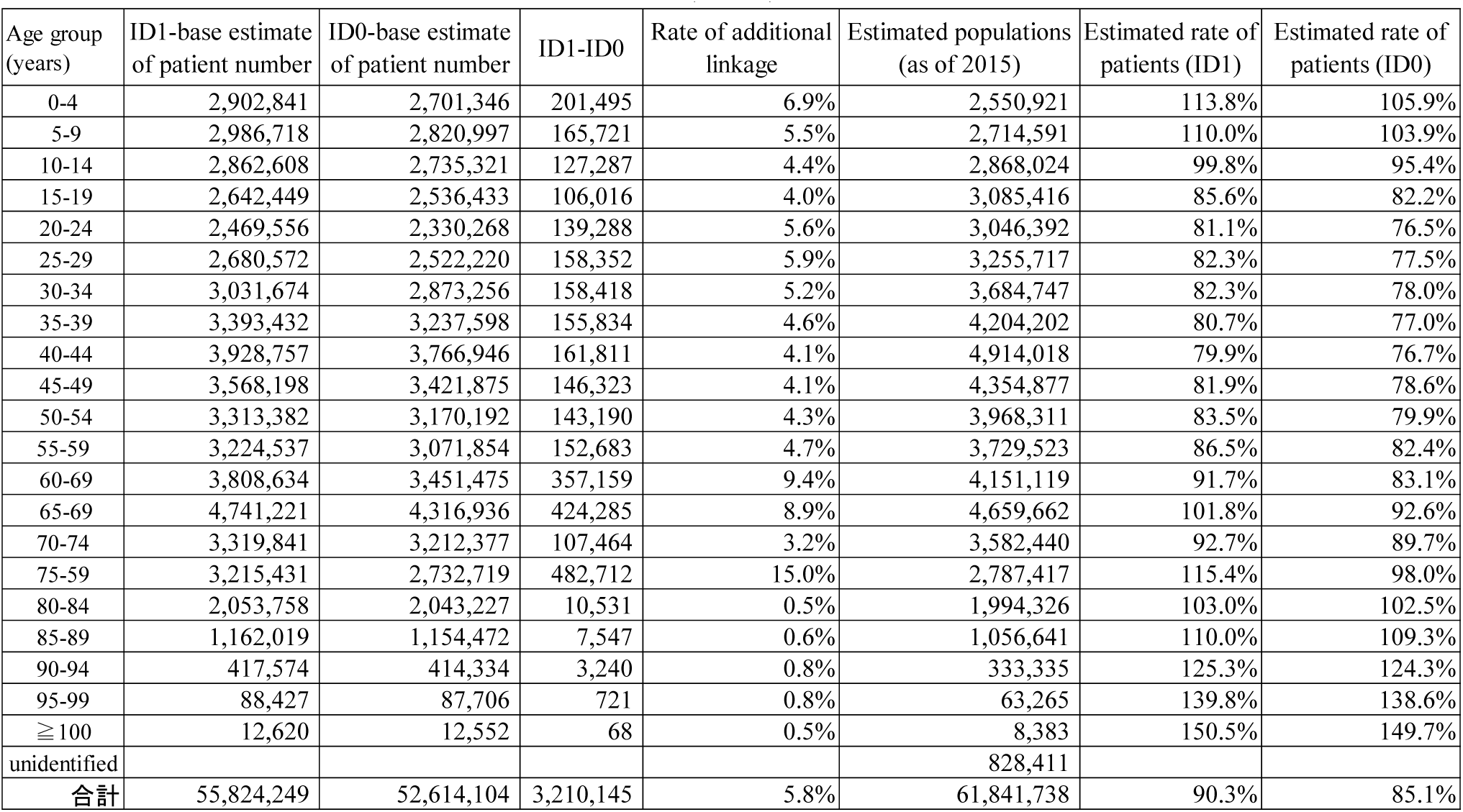
ID0- and ID1-based estimation of the rate of patients by age group in FY2015 (men)

| Age group (years) | ID1-base estimate of patient number | ID0-base estimate of patient number | ID1-ID0 | Rate of additional linkage | Estimated populations (as of 2015) | Estimated rate of patients (ID1) | Estimated rate of patients (ID0) |
| --- | --- | --- | --- | --- | --- | --- | --- |
| 0-4 | 2,902,841 | 2,701,346 | 201,495 | 6.9% | 2,550,921 | 113.8% | 105.9% |
| 5-9 | 2,986,718 | 2,820,997 | 165,721 | 5.5% | 2,714,591 | 110.0% | 103.9% |
| 10-14 | 2,862,608 | 2,735,321 | 127,287 | 4.4% | 2,868,024 | 99.8% | 95.4% |
| 15-19 | 2,642,449 | 2,536,433 | 106,016 | 4.0% | 3,085,416 | 85.6% | 82.2% |
| 20-24 | 2,469,556 | 2,330,268 | 139,288 | 5.6% | 3,046,392 | 81.1% | 76.5% |
| 25-29 | 2,680,572 | 2,522,220 | 158,352 | 5.9% | 3,255,717 | 82.3% | 77.5% |
| 30-34 | 3,031,674 | 2,873,256 | 158,418 | 5.2% | 3,684,747 | 82.3% | 78.0% |
| 35-39 | 3,393,432 | 3,237,598 | 155,834 | 4.6% | 4,204,202 | 80.7% | 77.0% |
| 40-44 | 3,928,757 | 3,766,946 | 161,811 | 4.1% | 4,914,018 | 79.9% | 76.7% |
| 45-49 | 3,568,198 | 3,421,875 | 146,323 | 4.1% | 4,354,877 | 81.9% | 78.6% |
| 50-54 | 3,313,382 | 3,170,192 | 143,190 | 4.3% | 3,968,311 | 83.5% | 79.9% |
| 55-59 | 3,224,537 | 3,071,854 | 152,683 | 4.7% | 3,729,523 | 86.5% | 82.4% |
| 60-69 | 3,808,634 | 3,451,475 | 357,159 | 9.4% | 4,151,119 | 91.7% | 83.1% |
| 65-69 | 4,741,221 | 4,316,936 | 424,285 | 8.9% | 4,659,662 | 101.8% | 92.6% |
| 70-74 | 3,319,841 | 3,212,377 | 107,464 | 3.2% | 3,582,440 | 92.7% | 89.7% |
| 75-59 | 3,215,431 | 2,732,719 | 482,712 | 15.0% | 2,787,417 | 115.4% | 98.0% |
| 80-84 | 2,053,758 | 2,043,227 | 10,531 | 0.5% | 1,994,326 | 103.0% | 102.5% |
| 85-89 | 1,162,019 | 1,154,472 | 7,547 | 0.6% | 1,056,641 | 110.0% | 109.3% |
| 90-94 | 417,574 | 414,334 | 3,240 | 0.8% | 333,335 | 125.3% | 124.3% |
| 95-99 | 88,427 | 87,706 | 721 | 0.8% | 63,265 | 139.8% | 138.6% |
| ≥ 100 | 12,620 | 12,552 | 68 | 0.5% | 8,383 | 150.5% | 149.7% |
| unidentified |  |  |  |  | 828,411 |  |  |
| 合計 | 55,824,249 | 52,614,104 | 3,210,145 | 5.8% | 61,841,738 | 90.3% | 85.1% |

**Table 3.** ID0- and ID1-based estimation of the rate of patients by age group in FY2015 (women)

| Age group (years) | ID1-base estimate of patient number | ID0-base estimate of patient number | ID1-ID0 | Rate of additional linkage | Estimated populations (as of 2015) | Estimated rate of patients (ID1) | Estimated rate of patients (ID0) |
| --- | --- | --- | --- | --- | --- | --- | --- |
| 0-4 | 2,748,574 | 2,562,889 | 185,685 | 6.8% | 2,436,785 | 112.8% | 105.2% |
| 5-9 | 2,819,348 | 2,667,262 | 152,086 | 5.4% | 2,585,196 | 109.1% | 103.2% |
| 10-14 | 2,679,450 | 2,563,528 | 115,922 | 4.3% | 2,731,293 | 98.1% | 93.9% |
| 15-19 | 2,679,371 | 2,561,397 | 117,974 | 4.4% | 2,922,972 | 91.7% | 87.6% |
| 20-24 | 3,032,249 | 2,779,923 | 252,326 | 8.3% | 2,921,735 | 103.8% | 95.1% |
| 25-29 | 3,413,184 | 3,096,815 | 316,369 | 9.3% | 3,153,895 | 108.2% | 98.2% |
| 30-34 | 3,794,206 | 3,489,924 | 304,282 | 8.0% | 3,606,131 | 105.2% | 96.8% |
| 35-39 | 4,059,079 | 3,784,893 | 274,186 | 6.8% | 4,111,955 | 98.7% | 92.0% |
| 40-44 | 4,550,591 | 4,280,884 | 269,707 | 5.9% | 4,818,200 | 94.4% | 88.8% |
| 45-49 | 4,095,073 | 3,858,235 | 236,838 | 5.8% | 4,307,927 | 95.1% | 89.6% |
| 50-54 | 3,754,610 | 3,532,266 | 222,344 | 5.9% | 3,961,985 | 94.8% | 89.2% |
| 55-59 | 3,620,039 | 3,372,870 | 247,169 | 6.8% | 3,785,723 | 95.6% | 89.1% |
| 60-69 | 4,141,780 | 3,782,352 | 359,428 | 8.7% | 4,303,891 | 96.2% | 87.9% |
| 65-69 | 5,137,773 | 4,787,259 | 350,514 | 6.8% | 4,984,205 | 103.1% | 96.0% |
| 70-74 | 3,905,513 | 3,765,333 | 140,180 | 3.6% | 4,113,371 | 94.9% | 91.5% |
| 75-79 | 4,002,735 | 3,409,676 | 593,059 | 14.8% | 3,489,439 | 114.7% | 97.7% |
| 80-84 | 2,965,788 | 2,944,547 | 21,241 | 0.7% | 2,967,094 | 100.0% | 99.2% |
| 85-89 | 2,135,719 | 2,116,096 | 19,623 | 0.9% | 2,060,616 | 103.6% | 102.7% |
| 90-94 | 1,142,237 | 1,130,903 | 11,334 | 1.0% | 1,015,785 | 112.4% | 111.3% |
| 95-99 | 374,850 | 371,411 | 3,439 | 0.9% | 296,082 | 126.6% | 125.4% |
| ≥ 100 | 76,207 | 75,684 | 523 | 0.7% | 53,380 | 142.8% | 141.8% |
| unidentified |  |  |  |  | 625,347 |  |  |
| Total | 65,128,376 | 60,934,147 | 4,194,229 | 6.4% | 65,253,007 | 99.8% | 93.4% |

### 3.3 ID track rate in a three-year period

The rates of the numbers of ID0 in FY2014 and FY2015 versus the number of ID0 in 2013 (ID track rates) were calculated (Tables 4 and 5). The ID track rate dropped every year by roughly 10%, although inclusion of cases of deaths and of no medical claim made must be noted. The non-track rate peaked in individuals aged 20-24 years, gradually declining in older age groups, and then increasing again in those aged ≥85 years.

**Table 4.** ID track rate between 2013 and 2015 (men)

| Age group (years) | No. of tracked patients |  |  | Non-track rate |  |
| --- | --- | --- | --- | --- | --- |
|  | 2013 | 2014 | 2015 | 2014 | 2015 |
| 0-4 | 2,771,836 | 2,605,080 | 2,492,733 | 6% | 10% |
| 5-9 | 2,817,459 | 2,645,331 | 2,502,248 | 6% | 11% |
| 10-14 | 2,793,074 | 2,593,271 | 2,376,648 | 7% | 15% |
| 15-19 | 2,493,326 | 2,138,046 | 1,846,311 | 14% | 26% |
| 20-24 | 2,291,451 | 1,783,726 | 1,545,973 | 22% | 33% |
| 25-29 | 2,612,312 | 2,125,291 | 1,856,644 | 19% | 29% |
| 30-34 | 2,916,629 | 2,462,412 | 2,177,310 | 16% | 25% |
| 35-39 | 3,388,503 | 2,931,083 | 2,606,639 | 13% | 23% |
| 40-44 | 3,624,797 | 3,168,144 | 2,832,335 | 13% | 22% |
| 45-49 | 3,183,234 | 2,822,016 | 2,558,981 | 11% | 20% |
| 50-54 | 2,982,234 | 2,687,216 | 2,475,269 | 10% | 17% |
| 55-59 | 3,048,311 | 2,772,717 | 2,575,662 | 9% | 16% |
| 60-69 | 3,832,411 | 3,436,340 | 3,210,298 | 10% | 16% |
| 65-69 | 3,747,865 | 3,480,790 | 3,321,085 | 7% | 11% |
| 70-74 | 3,319,978 | 3,136,271 | 2,979,843 | 6% | 10% |
| 75-79 | 2,642,922 | 2,481,235 | 2,359,388 | 6% | 11% |
| 80-84 | 1,905,220 | 1,761,457 | 1,621,891 | 8% | 15% |
| 85-89 | 1,054,270 | 925,399 | 805,457 | 12% | 24% |
| 90-94 | 343,818 | 278,434 | 222,183 | 19% | 35% |
| 95-99 | 78,689 | 55,938 | 39,185 | 29% | 50% |
| ≥100 | 11,215 | 6,823 | 4,100 | 39% | 63% |
| Total | 51,859,554 | 46,297,020 | 42,410,183 | 11% | 18% |

**Table 5.** ID track rate between 2013 and 2015 (women)

| Age group (years) | No. of tracked patients |  |  | Non-track rate |  |
| --- | --- | --- | --- | --- | --- |
|  | 2013 | 2014 | 2015 | 2014 | 2015 |
| 0-4 | 2,631,496 | 2,455,561 | 2,342,469 | 7% | 11% |
| 5-9 | 2,667,387 | 2,488,503 | 2,337,878 | 7% | 12% |
| 10-14 | 2,606,693 | 2,406,947 | 2,211,559 | 8% | 15% |
| 15-19 | 2,545,157 | 2,266,765 | 2,043,680 | 11% | 20% |
| 20-24 | 2,815,416 | 2,300,540 | 2,028,828 | 18% | 28% |
| 25-29 | 3,295,039 | 2,759,453 | 2,443,648 | 16% | 26% |
| 30-34 | 3,613,670 | 3,148,750 | 2,847,355 | 13% | 21% |
| 35-39 | 4,014,149 | 3,568,381 | 3,253,591 | 11% | 19% |
| 40-44 | 4,180,832 | 3,738,287 | 3,421,031 | 11% | 18% |
| 45-49 | 3,626,668 | 3,277,676 | 3,027,346 | 10% | 17% |
| 50-54 | 3,365,422 | 3,073,068 | 2,861,927 | 9% | 15% |
| 55-59 | 3,397,284 | 3,095,614 | 2,891,806 | 9% | 15% |
| 60-69 | 4,247,141 | 3,884,053 | 3,661,480 | 9% | 14% |
| 65-69 | 4,185,212 | 3,959,582 | 3,813,201 | 5% | 9% |
| 70-74 | 3,901,843 | 3,730,497 | 3,575,450 | 4% | 8% |
| 75-79 | 3,340,996 | 3,191,305 | 3,097,005 | 4% | 7% |
| 80-84 | 2,807,870 | 2,680,704 | 2,553,906 | 5% | 9% |
| 85-89 | 1,983,260 | 1,830,867 | 1,677,370 | 8% | 15% |
| 90-94 | 1,016,985 | 878,040 | 747,606 | 14% | 26% |
| 95-99 | 322,954 | 250,191 | 190,097 | 23% | 41% |
| ≥100 | 67,295 | 44,162 | 28,454 | 34% | 58% |
| Total | 60,634,782 | 55,030,960 | 51,057,702 | 9% | 16% |

Seven patterns of the yearly presence and/or absence of claims in a 3-year period were summarized together with ID1-based and ID0-based estimates of the patient numbers (Table 6). Track rates and non-track rates in a 3-year period are shown in Table 7.

**Table 6.** Patterns of the presence/absence of claims and the estimated patient number (ID0)

| Pattern | FY2013 | FY2014 | FY2015 | Estimated patient number (ID1) | Estimated patient number (ID0) |
| --- | --- | --- | --- | --- | --- |
| 1 | A | E | I | 77,592,278 | 89,087,842 |
| 2 | B | F |  | 16,449,344 | 8,975,145 |
| 3 | C |  | J | 4,312,101 | 5,264,523 |
| 4 | D |  |  | 21,515,184 | 9,818,854 |
| 5 |  | G | K | 17,239,198 | 10,204,568 |
| 6 |  | H |  | 8,819,987 | 4,996,498 |
| 7 |  |  | L | 21,809,048 | 8,935,644 |
| Total(ID1) | 119,868,907 | 120,100,807 | 120,952,625 |  |  |
| Total(ID0) | 113,146,364 | 113,264,053 | 113,492,577 |  |  |

**Table 7.** Patient track rates and non-track rates with ID1- and ID0-based approaches

| ID | FY | Estimated no. of patients | No. of patients to be tracked | No. of tracked patients (pre-adjustment) | Track rate (pre-adjustment) | Non-track rate (pre-adjustment) | Deaths per year | No claim*1 | A single year claim *2 | No. of tracked patients (post-adjustment)*3 | Track rate (post-adjustment) | Non-track rate (post-adjustment) |
| --- | --- | --- | --- | --- | --- | --- | --- | --- | --- | --- | --- | --- |
| ID1 | 2013 | 119,868,907 | 119,868,907 | 119,868,907 | 100.0% | 0.0% |  |  |  | 119,868,907 | 100.0% | 0.0% |
|  | 2014 | 120,100,807 | 119,868,907 | 94,041,622 | 78.5% | 21.5% | 1,256,359 | 4,312,101 | 8,819,987 | 108,430,069 | 90.3% | 9.7% |
|  | 2015 | 120,952,625 | 119,868,907 | 81,904,379 | 68.3% | 31.7% | 1,269,000 | 4,320,443 | 8,819,987 | 96,313,809 | 79.6% | 20.4% |
| ID0 | 2013 | 113,146,364 | 113,146,364 | 113,146,364 | 100.0% | 0.0% |  |  |  | 113,146,364 | 100.0% | 0.0% |
|  | 2014 | 113,264,053 | 113,146,364 | 98,062,987 | 86.7% | 13.3% | 1,269,000 | 5,264,523 | 4,996,498 | 109,593,008 | 96.8% | 3.2% |
|  | 2015 | 113,492,577 | 113,146,364 | 94,352,365 | 83.4% | 16.6% | 1,302,000 | 5,269,999 | 4,996,498 | 105,920,862 | 93.3% | 6.7% |

\*1 The number of patients who happened not to visit a medical institution.
\*2. The number of individuals who were healthy in the first year, and visited a medical institute in a following single year.
\*3. Number of tracked patients (post-adjustment)=Number of tracked patients (pre-adjustment)+ annual death number
+ Number of patients with no claim+ Number of patients with a single year claim

For example, the number of patients who visited a hospital in FY2013 is equal to A+B+C+D. Of those, the number of tracked patients in FY2014 is E+F. Therefore, the track rate of FY2014 patients in 2013 is (E+F)/(A+B+C+D). Similarly, the number of tracked patients in FY2015 is I+J, and the track rate in 2013 is (I+J)/(A+B+C+D).

These track rates are the rates of IDs entered in the database in certain years to those entered in the reference year, but not actual patient track rates because the absence of a claim in a certain year due to the following reasons were not taken into account: (1) death (≥ 1 million deaths per year); and (2) no insurance claim made (because patients did not visit hospitals/clinics). Thus, we adjusted the raw ID track rates with the above to estimate the actual patient track rates.

First, in pattern 3, claims were absent in 2014, but present in 2015, indicating that ID-linkage was not lost. Thus, the patient number in cell “J” in Table 6 was added to the number of tracked patients in 2014, and the corresponding estimate in 2016 obtained based on the total patient number was added to the number of tracked patients in 2015. Secondly, the patient number in cell “H” was added to the number of tracked patients in 2014 and 2015, because claims were made in a single year as intended (e.g. claims corresponding to basically healthy beneficiaries). Lastly, the annual number of deaths was added to the tracked patient number. With these adjustments, the ID1-based non-track rate dropped to 31.7%, while the ID0-based non-track rate dropped to 6.7%.

## 4. Discussion

### 4.1 Evaluation of ID0-based data linkage

This study proposed ID0, which is ID1 after a process wherein multiple ID1s corresponding to a given beneficiary are paired using ID2 and health care outcome in order to justify the linking of different ID1s, and the preceding ID1 is sequentially replaced by a newly appearing ID1. This approach is novel mainly because outcome information, in addition to ID1 and ID2, in a 3-year period is used to obtain a new variable ID0. The same ID2 will be assigned to different beneficiaries if they are of same sex, with identical date of birth and name. Given that assignment of the same ID1 to multiple beneficiaries occurs less frequently (except dependent twins of same sex supported by the same insured individual), ID1 was solely used in many statistical analyses of the NDB, and changes in ID1 during a given study period were neglected in such analyses. Even though both ID1 and ID2 were used for data aggregation, there were some issues where the data of different beneficiaries were linked even though death was noted in the outcome section, and data of the same beneficiaries were not linked because of a delayed claim due to loss of eligibility of insured individuals. The ID0-based approach, as newly proposed in this study, overcomes these problems.

An old ID1 and a new ID1 can coexist after loss of eligibility for the old insurance coverage. To increase data aggregation accuracy as much as possible, linkable data were searched using ID2 month by month, not at one time but for a period of 3 months after change in ID1. Claims associated with the same beneficiaries can be issued twice in 1 month due to changes in ID1 and/or ID2. This study showed that, even when both ID1 and ID2 were changed (quitting a job upon marriage, becoming independent beneficiaries upon divorce) at the same time, the old ID2 was occasionally used for a while, enabling data linking especially when claims were issued continuously (e.g. inpatient medical care and continuous outpatient treatment).

As a result, the ID0-based approach reduced the estimated number of male patients by 6.2%, and that of female patients by 7.1% compared with the corresponding ID1-based figures. The number of patients (but not total number of visits) exceeded the population estimates in several age groups, posing a significant concern with the ID1-based approach. The ID0-based approach appeared to yield a number closer to the actual patient number, except in some age groups (boys and girls aged 0-9 years, men aged ≥80 years, women aged ≥85 years), suggesting that this method is not perfect. It is noteworthy that the patient number here is the number of individual IDs in the NDB, and thus IDs of beneficiaries who died before the reference date for population statistics (usually October 1 in Japan) were counted. Also, changes in insurer due to finding a job, changing jobs, or retirement tend to peak in March at the end of the fiscal year. The fundamental solution to the current problems in ID-based data aggregation is assignment of unique lifetime ID to patients.

The rates of ID1s linked to only one ID2 were higher in medical inpatient claims and DPC claims than in medical outpatient claims. Inpatients (medical and DPC inpatients) are less likely to visit multiple medical institutions than outpatients, consequently, there would be less ID2 variations due to orthographic variations, which may explain such differences in ID1-ID2 pairing.

NDB data in a 3-year period were aggregated in this study. It must be noted that a certain number of patients became untrackable. Because ID0-based data linking is not perfect, and deaths cannot be identified clearly, the true patient trackability remains unknown. Even through deaths noted in the outcome section were taken into account, the track rate was as low as 70%. Improvement of accuracy of outcome information is anticipated.

Some age groups showed an estimated rate of patients of 100% or over after ID0-based data aggregation. This is clearly due to defects in the process of obtaining ID0, but no concrete explanation is currently available.

The Ministry of Internal Affairs and Communications estimates the population based on the latest national population census report, but the rate of the difference in figures between the national population census and the basic resident register to the figure in the basic resident register is approximately 1% (greater than 100 people)^6^. Also, this study used the 2013 mid-year population, not a population obtained after aggregation of a year’s worth of data. Errors may exist in population estimates, but problems in this ID0-based system need to be addressed in the future. ID1-based data aggregation estimated the patient number much greater than the population estimates, and 13% more than that estimated by ID0-based data aggregation, in the group aged 75-79 years. This may be mainly because everybody receives a new ID1 upon an insurer switch to the Health Insurance System for Latter-stage Elderly at age 75 years; ID1-based data aggregation is likely to process such newly assigned ID1s as those assigned to different individuals. Given that the majority of NDB data correspond to the elderly population, there is a risk of large discrepancies between estimates and actual patient numbers.

One of the problems of ID0-based data linking, on which we are currently working, lies in male-female differences. The rate of the patient number (number of individual IDs) to the population estimate was lowest in men aged 20-39 years (slightly higher than 70%) than in any other age groups. This agrees with the notion of a low hospital visiting rate among young and middle-aged adults. However, the rate of the patient number to the population estimate in women aged 20-39 years reached 90%, which is higher than those in adjacent age groups. There are two possible reasons. One is health insurance claims related to childbirth. Childbirth costs are self-covered in principle. Thus, the NDB does not contain related claims, except those for perinatal medical care for abnormal labor (e.g. forceps delivery, vacuum extraction, and Caesarean section). These exceptional claims may contribute to a high estimated rate of beneficiaries who required medical care in women aged 20-39 years. The other reason is that the name and the insurer often change in women in such age groups. The rate of women who changed their second name to their husbands’ second name was 96.2% in 2013, and consequently, ID2s of many women change after marriage^7^. If these women become dependents and are covered by their husbands’ insurers (i.e. change in insurer) their ID1s also change. Changes in name and insurer can occur in both men and women, but particularly more frequently in women in their twenties and thirties. If such women visit medical institutions during the period when their ID1s and ID2s are changing, claims issued are likely to be dealt with as those corresponding to different women.

Medical care continuity check, based on the location where medical care was delivered and diagnostic outcomes, may increase the accuracy of ID0-based data linking. If claims with the same ID indicate that visits to different medical institutions, distantly located, were made for completely different health conditions, it is appropriate to think that those claims correspond to different beneficiaries. Conversely, if claims with different IDs indicate the same location, age group, diagnosis, and therapy contents, they are more likely to correspond to the same beneficiary. Of course, some may frequently go to a different location on business and visit a medical institution there, and different individuals may receive the roughly same treatment in the same location. More precise data linking using other pieces of information, such as location, age group, and diagnosis, would be achievable in limited cases. But even so, many problems need to be solved to reduce the risk of type I errors. Nevertheless, ID0-based data aggregation is designed to eliminate type I errors, and thus claims with unlinked IDs are dealt with as those corresponding to different beneficiaries even though they are highly likely to correspond to the same beneficiary.

### 4.2 Rationale for using ID0

This study estimated the number of patients after ID0-based data linking. The accuracy of ID1 is extremely low, and thus ID1-based cohort studies may not yield valid outcomes. As described earlier, ID1 is a hash value generated from the insurer’s ID, and the beneficiary’s ID, data of birth, and sex, and changes in any of these elements result in changes in ID1. Indeed, ID1s of approximately 8 million beneficiaries change every year. Given that job change, transfer, and switching to the Health Insurance System for Latter-stage Elderly People occurs at any time of the year, it is advisable that the ID1-based approach be avoided even when analyzing NDB data of a single year. Use of both ID1 and ID2 is essential, but we did not opt for a less stringent approach, wherein data with at least one matched ID (either ID1 or ID2) were dealt with as those corresponding to the same beneficiary, to avoid linking data of different beneficiaries (e.g. twins and those with the same name). Our primary focus in this study is avoiding the linkage of claims belonging to different beneficiaries (type I errors).

It is noteworthy that certain conditions are required to obtain a valid ID0, as proposed by this study. First, this study used data of four types of health insurance claims (medical inpatient, medical outpatient, DPC, and pharmacy claims) in a 3-year period. In other words, these claims are those extractable from the NDB, regardless of the subgroup of patients (by special extraction). Thus, use of claims belonging to certain subgroups of patients may result in decreases in rate of ID pairing and reproducibility. The rate of ID0-based ID pairing will increase as more claims are used. Also, claims issued in a 1-month or longer period must be used.

An alternative ID currently under consideration is ID3. ID3 addresses orthographic variations (full- or half-width, with/without an additional zero in front) that cause unsuccessful linkage between data of specific health checkups and data of health insurance claims. ID3 was currently assigned to claims issued in 2015 and beyond, and its use will be extended to older claims. ID3 is a more precise version of ID1, and the accuracy of data linking will increase when ID2 is used in combination with ID3.

Dental claims were not used in this study, but the approach shown in this study is applicable, by merging an intermediate table made from data of medical inpatient care, medical outpatient care, DPC and dental care claims, with an intermediate table made from pharmacy claims.

An intermediate table made from data of medical inpatient care, medical outpatient care, and DPC claims was merged with that made from data of pharmacy claims in this study, to increase the ID-linkage rate. The rationale behind this is that medical care claims and associated pharmacy claims were likely to be issued in the same period and processing these two types of claims at the same time will reduce one-to-one ID pairing/linking. This must be avoided, being the exact point that we focused on, and a critical modification was made in this study.

### 4.3 Limits of the NDB

The Japanese public health insurance system basically allows insured medical care, but not a mixed medical case series (combination of insured and uninsured treatment performed in the same series of medical treatment). The NDB is a complete enumeration of insured treatment claims, and thus does not contain information associated with uninsured treatment (e.g. some advanced therapies, esthetic treatment, immunization, and health checks), publicly funded health care (e.g. provided to those who receiving income support), and diagnostic procedures. This fact needs to be considered when interpreting the outcomes of NDB analyses.

Also, the fact that the NDB is a complete enumeration of insured treatment claims, without inclusion of information associated with uninsured treatment and publicly funded health care must be clearly stated for appropriate discussion on differences between completely insured treatment and actual care provided.

Furthermore, it must be noted that names of health conditions in the NDB include undetermined diagnoses, and predetermined names of conditions that must be mentioned on the medical insurance claims for remuneration. In other words, names of conditions in the NDB are not necessarily confirmed diagnoses. When using the NDB, confirmed diagnoses need to be distinguished from the above temporarily assigned names of conditions, thereby determining the true names of health conditions in given individuals.

## 5. Conclusions

This study proposed a new innovative personal ID (ID0) for analysis of the NDB and evaluated ID0-based data aggregation that addressed the efficiency of existing data aggregation methods. Compared with existing methods using either ID1 or a combination of ID1 and ID2, the ID0-based method showed higher accuracy in data linking. However, problems where estimated patient number exceeded the actual population sizes remained in the following subgroups: children; old-old adults; and women of reproductive age. Nevertheless, the ID0 is the best variable to link data corresponding to a given individual, and thus it is recommended to use ID0 instead of ID1 when estimating the patient number.

## Acknowledgement

This study was part of the 2016/2017 Health and Labour Sciences Research Grant (Research on Regional Medical) project entitled “Study on implementable measures required for: differentiation of hospital beds depending on purpose; cooperation between different purpose divisions; and efficient use of hospital beds” and the 2016 Japan Agency for Medical Research and Development, Cross-regional ICT utilization project entitled “Study on medical performance evaluation including pharmacoepidemiologic approaches using a large-volume of electronic medical care data including health insurance claims”.

